# Pelvic bone structures in free-ranging Eurasian lynx (*Lynx lynx*) from Switzerland: a radiological pelvimetry study

**DOI:** 10.1101/646935

**Authors:** Fanny Morend, Johann Lang, Beatriz Vidondo, Marie-Pierre Ryser-Degiorgis

## Abstract

The observation of severe pelvic malformations in Eurasian lynx (*Lynx lynx*) from a population reintroduced to Switzerland raised the question as to whether inbreeding may contribute to the development of congenital pelvic malformations. We aimed at providing baseline data on the pelvic morphology of Eurasian lynx from the reintroduced populations in Switzerland, at assessing potential differences in pelvic conformation between the two main Swiss populations, among age classes and between sexes, and at detecting pelvic anomalies. We performed measurements of 10 pelvic parameters on the radiographs of 57 lynx of both sexes and different ages taken from 1997-2015. We calculated two ratios (vertical diameter/acetabula; sagittal diameter/transversal diameter) and two areas (pelvic outlet and inlet) to describe the shape of the pelvis. Our results showed that the Eurasian lynx is a mesatipelvic species, with a pelvis length corresponding to approximatively 20% of the body length. We found no statistically significant differences between the two examined populations but observed growth-related pelvis size differences among age groups. Sexual dimorphism was obvious in the adult age group only: two parameters reflecting pelvic width were larger in females, likely to meet the physiological requirements of parturition. By contrast, pelvis length, conjugata vera, diagonal conjugata, vertical diameter and sagittal diameter were larger in males, in agreement with their larger body size. Accordingly, the ratio between the sagittal and transversal diameters was significantly larger in males, i.e. adult males have a different pelvic shape than adult females. Furthermore, pelvimetry highlighted one adult individual with values outside the calculated reference range, suggesting a possible congenital or developmental pathological morphology of the internal pelvis. Our work generated baseline data of the pelvic morphology including growth and sexual dimorphism of the Eurasian lynx. These data could also be useful for estimating age and sex in skeletal remains.

## Introduction

The Eurasian lynx (*Lynx lynx*) vanished from Switzerland and other European countries at the end of the 19th century. Following habitat improvementy and prey recovery, lynx reintroductions took place in the 1970s, with lynx imported from the Capathian Mountains in Slovakia. In Switzerland, two small distinct lynx populations resulted from these releases: one in the Swiss Alps extending slightly into France and Italy, and another one in the Jura Mountains at the border with France, both estimated at approximatively 100-150 independent lynx, with a total of approximately 130-140 lynx in the Swiss territory (1). However, these populations are now characterized by a low genetic variability and a high inbreeding coefficient (2).

A lynx health monitoring program has been conducted at the Centre for Fish and Wildlife Health (FIWI) in Switzerland for several decades, including necropsies of carcasses, clinical examinations of live lynx and systematic sampling. From 2004 onwards, there has been increasing concern about inbreeding depression, in particular in the Alpine population (3). Among other abnormalities, pelvic malformations were detected in two orphaned lynx. In 2006 a juvenile female brought to a rehabilitation station died of a peritonitis following a rectal perforation due to a foreign body, in conjunction with a severe colitis associated with *Yersinia tuberculosis.* The week before death, the animal had experienced an episode diarrhoea followed by tenesmus. Post-mortem examination revealed a severe pelvic malformation. The pelvic canal appeared abnormally narrow and the left hemipelvis was inwardly rotated (Fig1A, B). Subsequently, another case with a similar history was found in the archive of the FIWI, and the examination of the pelvis preserved at the Museum of Natural History of Bern revealed similar malformations (Fig 1C). This raised questions regarding the occurrence and potential impact of congenital pelvic malformations in lynx in Switzerland. Since then, pelvic radiographs have been systematically taken of dead lynx submitted to necropsy and of live lynx to be translocated for reintroduction programs in neighbouring countries. Obvious pelvic malformations have not been detected, but thorough analyses of radiographs were not performed.

**Fig 1.**
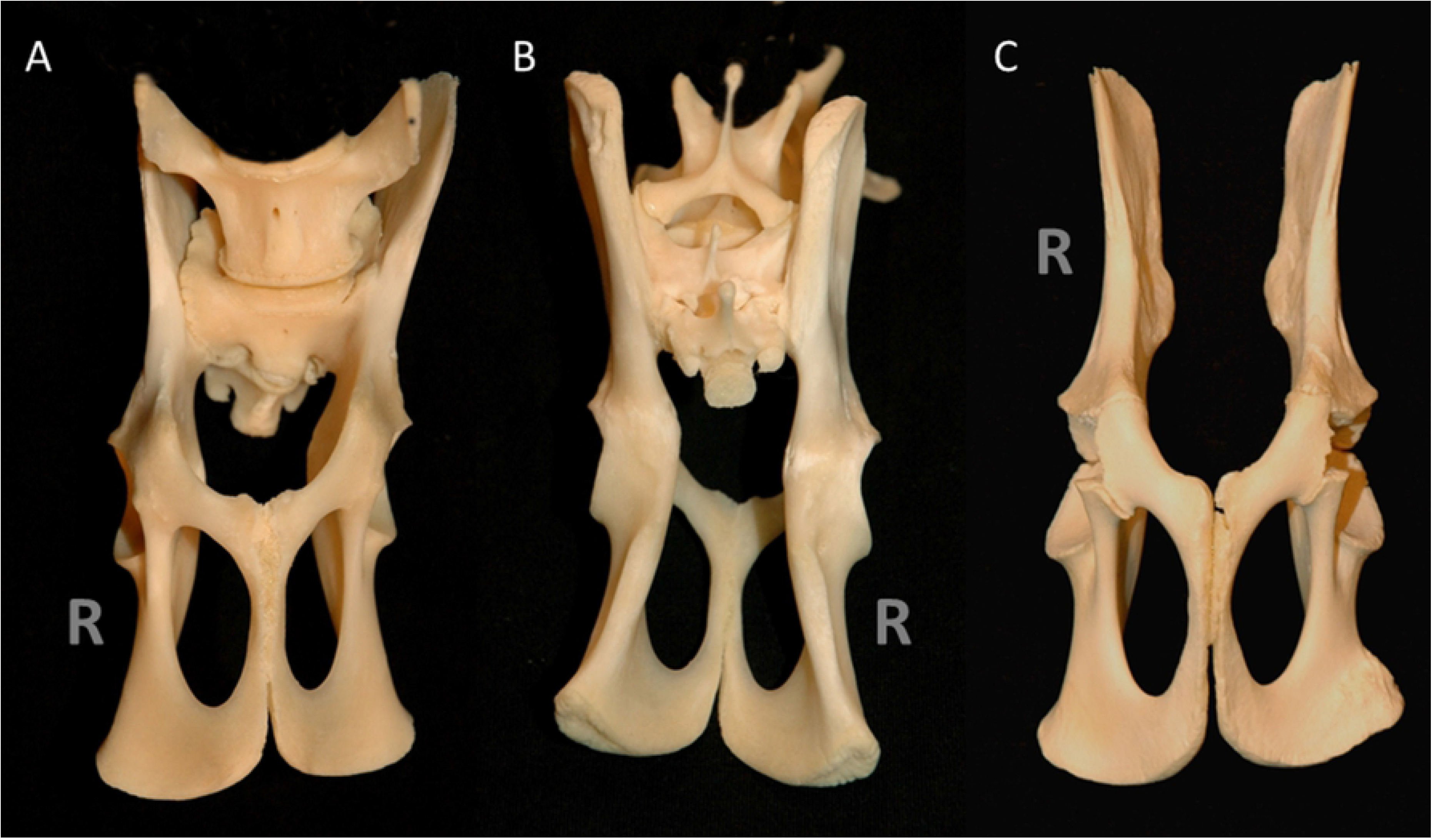
Abnormal pelvis, Eurasian lynx. (A) Ventral view and (B) Dorsal view, juvenile female, north-eastern Swiss Alps (specimen W06/1532/090, BEHTLI): The pelvis canal is abnormally narrow and the left hemipelvis is rotated towards the centre. (C) Juvenile male, north-western Alps (specimen W98/3505), ventral view (C): The left hemipelvis is slightly rotated towards the centre. R = right body side.

Narrowing of the pelvis or its deformation may occur due to malunion of an earlier fractured pelvis, metabolic disorders or congenital pelvic malformations, and it may result in complications such as obstipation and dystocia (4–6). Currently there are no studies proving the heritability of pelvic characteristics in felids. Nevertheless, it has been demonstrated in Boston terrier dogs that 26 per cent of the offspring’s pelvic shape could be explained by the pelvic shape of the parents (5,7,8). Thus, if pelvic shape were also inherited in lynx, pelvic malformation and narrowing may occur at an increased frequency in the inbred lynx population in Switzerland. This may have an impact on population dynamics in the long-term due to fatalities linked to chronic obstipation and dystocia.

Radiographic examination followed by pelvic measurements is a valuable method to assess pelvic narrowing or malformation. Pelvimetry consists of measuring distances and angles between pelvic structures, either by palpation or using radiographs. It can be done by palpation or on x-rays. Radiographic pelvimetry is often used to calculate the pelvic size in humans, and has also been employed in domestic animals such as cats (5,8), dogs (9,10), cattle (11), sheep (12) and Indian buffaloes (13).

Switzerland serves as a source for lynx reintroductions in other European countries and it is essential to thoroughly assess the health status of the populations. The objectives of the present study were: (i) to deliver baseline data on the pelvic morphology of Eurasian lynx from the reintroduced populations in Switzerland; (ii) to assess potential differences in pelvic conformation between these two populations, among age classes and between sexes; and (iii) to detect pelvic abnormalities that may have been missed by the routine evaluation of the radiographs of single cases.

Pelvimetry generated baseline data of the pelvic morphology, including growth and sexual dimorphism of the Eurasian lynx, and highlighted one adult individual with values outside the calculated reference range, suggesting a possible congenital or developmental pathological morphology of the internal pelvis.

## Material and Methods

### Ethics Statement

None of the animals concerned by this study was killed for research purposes. All of the samples originated either from free-ranging wildlife found dead in the field or from live animals captured within the framework of conservation actions. Collection and sampling of all animals respected Switzerland’s legislation (922.0 hunting law; and 455 animal protection law, including legislation on animal experimentation; www.admin.ch). Anesthesia, handling and examination of live lynx were carried out by experienced veterinarians after obtention of all required permits for capturing and handling lynx (for details see (14).

### Material

Radiographs of 100 lynx taken from 1997 to 2015 were available from the archives of the Vetsuisse Faculty of the University of Bern. Of these lynx, 86 were found dead and necropsied at the FIWI and 14 were captured in the field and taken to a quarantine station before translocation or rehabilitation, under veterinary supervision of the FIWI staff. Live lynx were trapped and anesthetized following established protocols as described elsewhere (15) and with all permits required by the Swiss legislation. Age was determined based on field data (animals identified as kittens) or tooth cementum analysis (Matson’s Laboratory, Manhattan, USA), or estimated using morphological criteria such as body size and weight, growth plates, tooth replacement and tooth wear. Age classes were defined as previously described (16): juveniles included lynx in their first year of life; subadults ranged from 12-24 months for females and from 12-36 months for males; and adults corresponded to females >24 months and males >36 months (post-growth period). Body length was measured from the tip of the nose to the base of the tail, following the back of the lynx lying in lateral recumbency (physiological position).

All dead animals were radiographed at the small animal hospital of the University of Bern. Fourteen live lynx were also radiographed at the hospital but for most of them it was done with a portable unit at the quarantine station. The x-ray units used to obtain the radiographs included a General Electrics Prestige SI (65kW), a Swissray Gen-X-80kW unit, a Siemens Polydoros LX Lite Power Up 100kW, and a portable Gierth HF 300 (Riesa, Germany) for radiographs taken at the quarantine station. The imaging systems included a Fuji FCR AC-3 and Fuji FCR Profect one reader. The imaging plates were FCR ST-V general purpose plates, standard dual side imaging plates (ST-BD) and dual sided mammographic HR-BD plates. Depending on the imaging system used, focus-film distance was 100 to 120 centimetres, and the exposition varied between 54 kV / 20 mAs to 70 kV / 25 mAs; and 63 kV / 5 mAs for the mobile system. Orthogonal radiographs of the pelvis were performed in the dorsal and either right or left lateral recumbency, with the aim of obtaining a symmetrical positioning of the pelvis.

### Pelvimetry

We excluded thirty-eight out of 100 lynx (38%) from the study because there was only one projection available, the pelvis was incompletely imaged, or because recent pelvic fractures did not allow exact measurements. We considered the radiographs of 62 lynx as being of sufficient quality for pelvimetry. We additionally conducted pelvimetry on radiographs of the juvenile female BEHTLI, in which a severe pelvic deformity had been observed (W06/1532/090; Fig 1A and 1B). Data collection for the second case with a similar history of pelvic malformation (juvenile male W98/3505; Fig 1C) was not possible, because radiographs were not available for this animal.

We performed all pelvic measurements directly on the radiographs with the radiology information system IMPAX EE (Agfa Health care, Bonn Germany), using a measurement tool with 0.01 cm accuracy. We took each measurement three times and retained the average of the three values. Similarly to previous studies in domestic cats (5,8), we made five measurements in the left-right or right-left lateral projection (Fig 2A) and five in the ventro-dorsal projection (Fig 2B). In addition, we calculated two ratios between the height and width of the pelvis: vertical diameter/acetabula (VD/AC) and sagittal diameter/transversal diameter (SGD/TD). Furthermore, we estimated the pelvic inlet area (PIA) and pelvic outlet area (POA) as follows: PIA = (CV/2 + TD/2)^2^ × π, and POA = (VD/2 + AC/2)^2^ × π (10), with CV corresponding to the distance between the promontory and cranial pubic symphysis (or conjugata vera, Fig 2A). The pelvic inlet area (PIA) is a planar surface which corresponds to the boundary between the pelvic and the abdominal cavity. The pelvic outlet area (POA) represents the lower circumference of the lesser pelvis.

**Fig 2.**
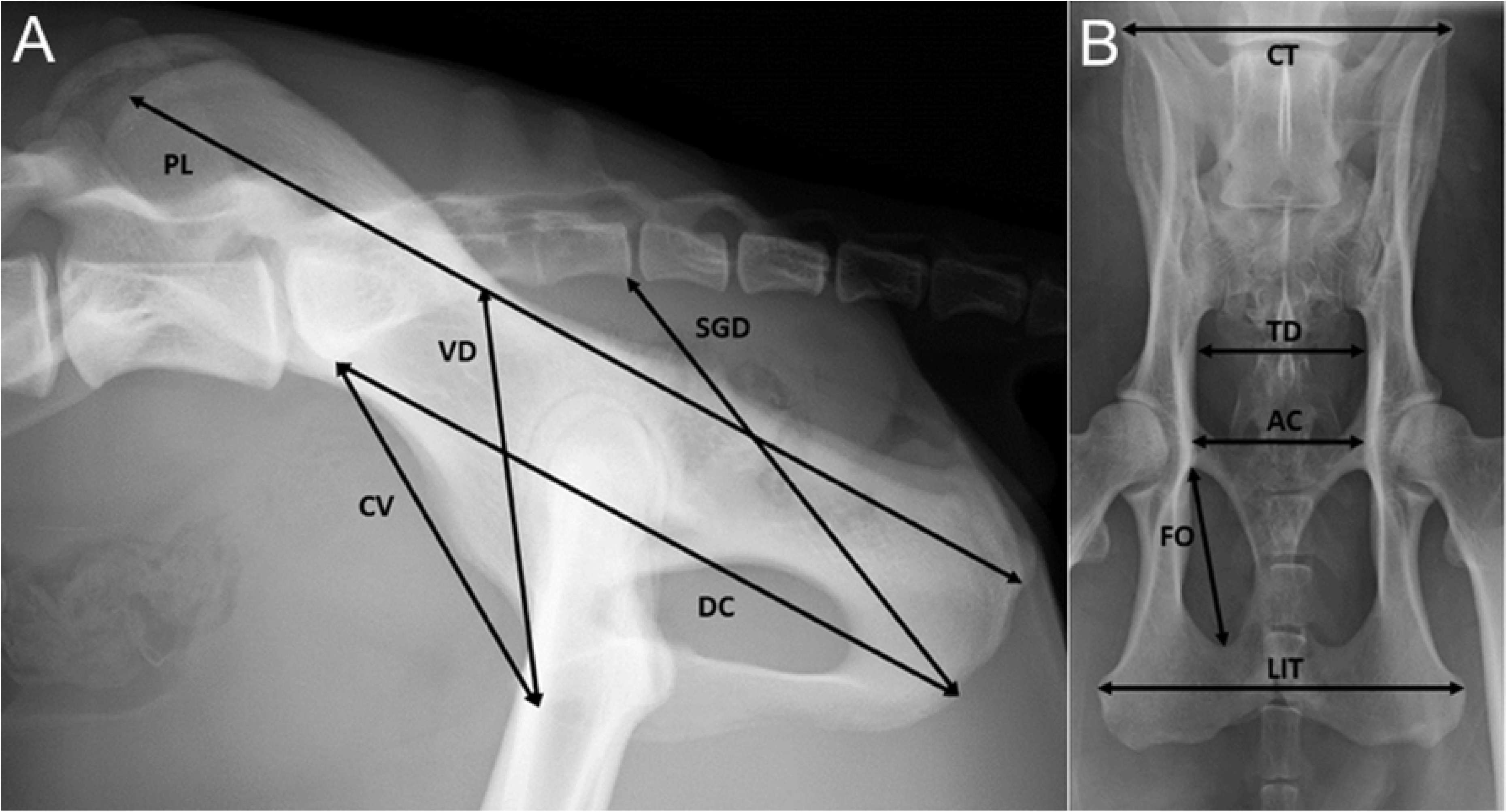
Measurements taken for pelvimetry. Pelvis, Eurasian lynx, subadult male (A) Left-right latero-lateral radiograph. Pelvis length (PL); conjugata vera (CV) = distance between the promontory and cranial pubic symphysis; diagonal conjugata (DC) = distance between the promontory and caudal pubic symphysis; vertical diameter (VD) = vertical distance between the cranial portion of the pubic symphysis and the mid sacrum; sagittal diameter (SGD) = distance between caudal sacral and caudal symphysis. (B) Ventro-dorsal radiograph. Inner length of the foramen obturatum (FO); coxal tuberosities (CT) = horizontal distance between the coxal tuberosities; transversal diameter (TD) = horizontal distance between ilial shafts; acetabula (AC) = horizontal distance between acetabula; lateral ischial tuberosities (LIT) = horizontal distance between the lateral ischial tuberosities.

### Data analysis and statistics

We stored the data in a MS Excel© spreadsheet and imported them into the statistical software NCSS (NCSS 10 Statistical Software, 2015; NCSS, LLC; Kaysville, Utah, USA, ncss.com/software/ncss). We excluded five <4 month-old lynx from the data analysis because their pelvic values were much lower (pelvis length <10 cm) than those of other juveniles and these animals were too few to create an additional age class. The 57 remaining lynx included 32 animals from the Alps, 23 from the Jura, and two from a recently reintroduced population nucleus in north-eastern Switzerland. There were 26 males, 30 females and one animal of unknown sex (adult, Jura population); and 24 juveniles (4-12 months old), 11 subadults and 22 adults.

We used the Shapiro-Wilk normality test to assess the normal distribution of each variable. First, we tested for differences between the two main populations (Alps and Jura), among age classes and between sexes for each variable (i.e. pelvic measurements) in a univariable approach, using either the parametric unpaired Student t test (if variables were normally distributed) or the Mann-Whitney U test (if variables were not normally distributed). Subsequently, population, age class and sex were combined in a multivariable approach. We applied a MANOVA for each pelvic measurement (outcome variable), including the population, age class and sex as explanatory variables. The level of significance was set to 0.05. In addition to the five juveniles <4 months, we excluded the adult lynx of unknown sex and the two lynx from the population in north-eastern Switzerland from the multivariable models, i.e., these models included 54 individuals: 32 from the Alps (16 juveniles, 4 subadults and 12 adults) and 22 from the Jura (7 juveniles, 6 subadults and 9 adults). Thus, the results apply to specimens >4 months from the two populations Alps and Jura.

Male Eurasian lynx are expected to be larger in size than females, at least in the subadult and adult age classes (17). Therefore, we assessed differences in body length and pelvis length between sexes for each of the three age classes using unpaired Student t tests. Additionally, we calculated the ratio between the pelvis length and the body length for each specimen, and we tested for differences between the ratio of males and females for each of the three age classes using unpaired Student t tests or Mann-Whitney tests as appropriate.

We did not include BETHLI in the statistical analyses but we discuss how this animal’s data compares to the measurements of six other specimens (three females and three males) of the same age class and with similar pelvic length (controls).

## Results

Descriptive statistics of the radiographic pelvic parameters per age class and sex are shown in Tables 1 and 2. We detected two outliers: an adult male from the Alps with low VD (specimen W11/4314/220, SIBO), and a juvenile male from the Alps with high CT and TD (W09/1096/058; Fig 3).

**Fig 3.**
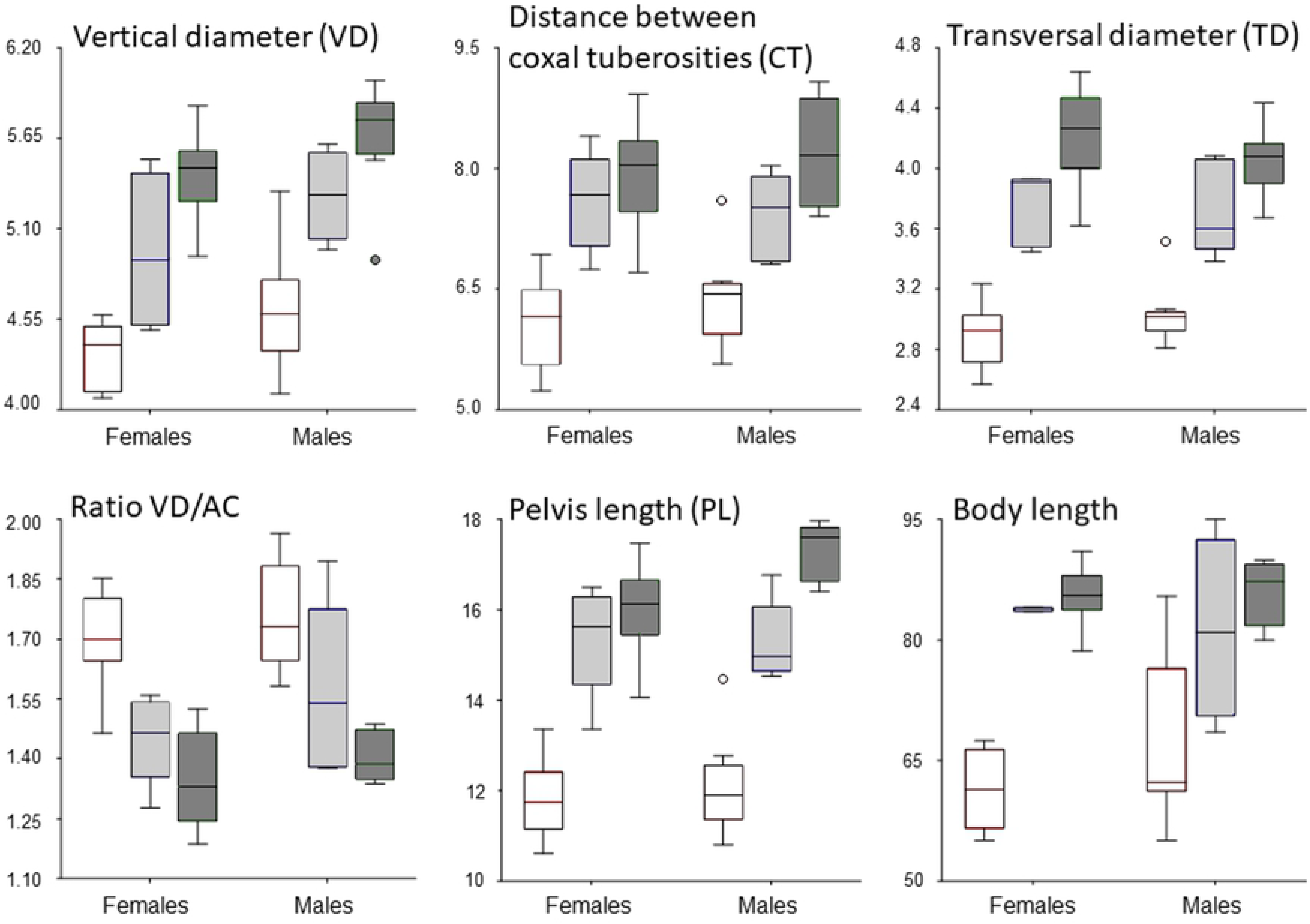
Box plots showing selected pelvic parameters and body length of Eurasian lynx. Longitudinal parameters and body length are given in centimeters. For each sex, data are indicated for juveniles (left, red), subadults (middle, blue) and adults (right, green). Outliers were detected for the vertical diameter of an adult male (low value); and for the horizontal distance between the coxal tuberosities, the transversal diameter and the length of the pelvis for a juvenile male (high values). The VD/AC ratio correspond to the vertical diameter divided by the distance between the acetabula.

**Table 1.** Radiographic parameters of female Eurasian lynx.

| Pelvic measurements<sup>a</sup> | Juveniles | Subadults | Adults |
| --- | --- | --- | --- |
| **Linear distances (cm)** |  |  |  |
| PL | 11.79±0.80<br>(10.60–13.37) | 15.38±1.21<br>(13.37–16.50) | 16.05±0.93<br>(14.07–17.47) |
| CV | 4.79±0.20<br>(4.52–5.09) | 5.53±0.32<br>(5.11–5.99) | 5.57±0.27<br>(5.09–5.92) |
| DC | 8.43±0.48<br>(7.89–9.39) | 10.56±0.54<br>(9.75–11.13) | 11.02±0.55<br>(9.75–11.7) |
| VD | 4.33±0.19<br>(4.07–4.58) | 4.96±0.47<br>(4.48–5.52) | 5.42±0.25<br>(4.93–5.85) |
| SGD | 5.69±0.33<br>(5.17–6.32) | 6.85±0.78<br>(5.49–7.38) | 7.53±0.45<br>(6.78–8.04) |
| FO | 3.12±0.27<br>(2.80–3.58) | 4.10±0.31<br>(3.78–4.46) | 4.29±0.31<br>(3.71–4.89) |
| CT | 6.09±0.53<br>(5.23–6.93) | 7.60±0.61<br>(6.75–8.40) | 7.93±0.60<br>(6.71–8.92) |
| TD | 2.89±0.19<br>(2.57–3.24) | 3.74±0.24<br>(3.45–3.93) | 4.22±0.31<br>(3.62–4.64) |
| AC | 2.55±0.24<br>(2.22–3.06) | 3.43±0.37<br>(2.94–3.77) | 4.04±0.35<br>(3.36–4.49) |
| LIT | 5.77±0.44<br>(5.23–6.50) | 7.69±0.67<br>(6.92–8.48) | 8.74±0.52<br>(7.57–9.26) |
| **Areas (cm<sup>2</sup>)** |  |  |  |
| PIA | 46.52±4.56<br>(39.44–54.15) | 67.67±7.32<br>(57.46–77.18) | 75.48±7.80<br>(59.58–84.84) |
| POA | 37.31±4.46<br>(31.04–44.65) | 55.72±10.28<br>(43.28–67.83) | 70.55±7.21<br>(54.06–79.17) |
| **Ratios (height/width)** |  |  |  |
| SGD/TD | 1.97±0.08<br>(1.85–2.18) | 1.87±0.14<br>(1.59–1.96) | 1.79±0.11<br>(1.58–1.93) |
| VD/AC | 1.70±0.11<br>(1.46–1.85) | 1.45±0.11<br>(1.28–1.56) | 1.35±0.11<br>(1.19–1.52) |

Mean, standard deviation and range (minimum-maximum) of the parameters of 30 free- ranging female Eurasian lynx (*Lynx lynx*) from Switzerland, 1997-2015.
<sup>a</sup> Pelvis length (PL), distance between the promontory and cranial pubic symphysis (CV), distance between the promontory and caudal pubic symphysis (DC), vertical distance between the cranial portion of the pubic symphysis and the mid sacrum (VD), distance between caudal sacral and caudal symphysis (SGD), inner length of the foramen obturatum (FO), horizontal distance between the coxal tuberosities (CT), transversal diameter (TD), horizontal distance
between acetabula (AC), horizontal distance between the lateral ischial tuberosities (LIT), pelvic inlet area (PIA), pelvic outlet area (POA).

**Table 2.** Radiographic parameters of male Eurasian lynx.

| Pelvic measurements<sup>a</sup> | Juveniles | Subadults | Adults |
| --- | --- | --- | --- |
| *Linear distances (cm)* |  |  |  |
| **PL** | 12.13±0.99<br>(10.80–14.47) | 15.29±0.85<br>(14.53–16.77) | 17.3±0.62<br>(16.40–17.97) |
| **CV** | 5.00±0.43<br>(4.25–5.84) | 5.54±0.30<br>(5.09–5.89) | 5.89±0.36<br>(5.21–6.42) |
| **DC** | 8.68±0.62<br>(7.40–10.33) | 10.61±0.84<br>(9.54–11.7) | 11.68±0.64<br>(10.5–12.43) |
| **VD** | 4.59±0.33<br>(4.10–5.33) | 5.30±0.27<br>(4.97–5.62) | 5.67±0.34<br>(4.91–6.03) |
| **SGD** | 5.93±0.52<br>(5.08–7.03) | 7.23±0.58<br>(6.77–8.15) | 8.05±0.47<br>(7.27–8.50) |
| **FO** | 3.28±0.33<br>(2.78–3.85) | 4.00±0.35<br>(3.62–4.43) | 4.44±0.32<br>(4.05–4.88) |
| **CT** | 6.37±0.53<br>(5.57–7.59) | 7.40±0.54<br>(6.81–8.03) | 8.18±0.65<br>(7.40–9.08) |
| TD | 3.02±0.18<br>(2.81–3.51) | 3.73±0.31<br>(3.38–4.08) | 4.04±0.22<br>(3.67–4.43) |
| AC | 2.62±0.17<br>(2.35–2.87) | 3.44±0.58<br>(2.69–4.06) | 4.01±0.27<br>(3.54–4.39) |
| LIT | 6.03±0.70<br>(4.89–7.30) | 7.79±0.81<br>(6.94–8.86) | 8.85±0.72<br>(7.77–9.94) |
| **Areas (cm<sup>2</sup>)** |  |  |  |
| PIA | 50.74±7.38<br>(41.40–68.71) | 67.64±7.56<br>(56.39–73.90) | 77.74±8.12<br>(61.93–87.20) |
| POA | 40.97±4.94<br>(34.70–52.72) | 60.43±11.38<br>(47.74–73.59) | 74.23±8.53<br>(56.08–83.81) |
| **Ratios (height/width)** |  |  |  |
| SGD/TD | 1.96±0.11<br>(1.76–2.13) | 1.94±0.07<br>(1.84–2.01) | 1.99±0.08<br>(1.88–2.13) |
| VD/AC | 1.76±0.14<br>(1.58–1.96) | 1.57±0.22<br>(1.38–1.89) | 1.40±0.06<br>(1.34–1.49) |

Mean, standard deviation and range (minimum-maximum) of the radiographic pelvic parameters of 26 free-ranging male Eurasian lynx (*Lynx lynx*) from Switzerland, 1997-2015.
<sup>a</sup> Pelvis length (PL), distance between the promontory and cranial pubic symphysis (CV), distance between the promontory and caudal pubic symphysis (DC), vertical distance between the cranial portion of the pubic symphysis and the mid sacrum (VD), distance between caudal sacral and caudal symphysis (SGD), inner length of the foramen obturatum (FO), horizontal distance between the coxal tuberosities (CT), transversal diameter (TD), horizontal distance between acetabula (AC), horizontal distance between the lateral ischial tuberosities (LIT), pelvic inlet area (PIA), pelvic outlet area (POA).

### Population

The mean of nearly all measured parameters was slightly larger in the Alps than in the Jura for all sex/age classes (Table A in S1 Table). Only the VD/AC ratio was significantly larger in adult lynx from the Jura (1.42 ± 0.09) than from the Alps (1.33 ± 0.09) when applying the unpaired Student t test (P=0.041) but when all explanatory variables were taken into account in the MANOVA, none of the pelvic parameters significantly differed between populations.

### Age

All linear radiographic pelvic measurements increased with age (Table 1 and 2, Fig 3). There was a statistically significant difference (MANOVA) between the three age groups in both males and females for all parameters except VD (Table B in S1 Table). This parameter also increased from the juvenile to the adult class and there was an outlier among adults (SIBO, 6 years old: VD=49.10 mm; range without this outlier: 55.17–60.03). Similarly, in the juvenile group one lynx (W09/1096/058) showed higher values (CT=75.93 mm; range without this outlier: 55.70–65.93; and TD=35.13 mm; range without this outlier: 28.07–30.67).

Regarding the pelvic height/width relationships, the VD/AC ratio significantly decreased with increasing age in both sexes (Table 1 and 2, Fig 3). The SGD/TD ratio did not significantly differ between age groups (Table 1 and Table 2).

### Sex

Females had generally lower mean values than males for the same measurement in all age classes. Among juveniles, a significant difference was found only for VD (Table B in S1 Table). Among subadults, differences between sexes were even less marked than in the juvenile age class and, by contrast, some parameters were lower in males than females (pelvis length, FO, CT); no statistically significant differences were detected for any of the measurements. Differences between sexes were most marked among adults, in which significant differences were present for the pelvis length (Fig 3), diagonal conjugata (DC), CV, VD, SGD and the SGD/TD ratio, all of them being characterized by higher values in males.

There were no statistically significant differences between sexes for the pelvic ratio VD/AC, whatever the age class. The pelvic height/width relationships did not differ between sexes among juveniles and subadults either, meaning that males and females of these two age groups have the same pelvic shape. Statistically significant differences were found only within the adult class, and only for the SGD/TD ratio, which was larger in males.

The MANOVA pointed at a significant influence of the combination of both age and sex for only one pelvic measurement: the pelvis length (Table A in S1 Table), which was larger in males in all age groups.

### Relationship between pelvic length and body length

The body size of the lynx included in this study and the ratio between the mean pelvis length and the mean body length are presented for all age/sex categories in Fig 3 and Table 3. Although the mean body length was higher in males than females among subadults and adults, these differences were not statistically significant (unpaired Student t test). Regarding the ratio pelvic/body length, there were no statistically significant differences (unpaired Student t test) between the sexes in any of the age classes, meaning that in lynx of any age the pelvic length is proportional to the body length, the pelvis corresponding to approximately 20% of the body length.

**Table 3.**
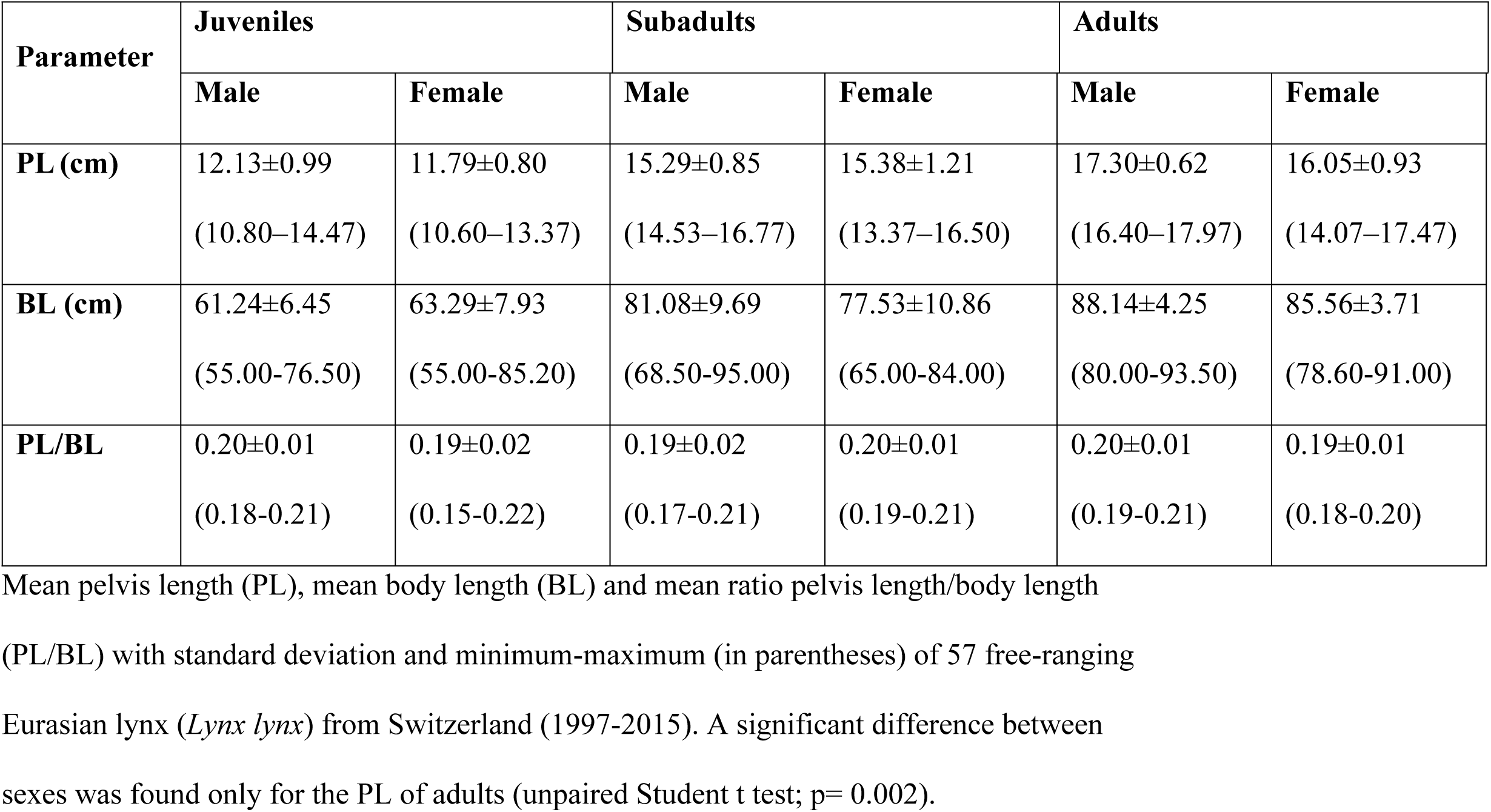
Pelvic parameters of Eurasian lynx considering sex and age class.

### Malformed juvenile pelvis

Nine out of 13 measures of BEHTLI’s pelvis (Table 4) were smaller than the minimum of the control specimens, including five longitudinal parameters (CV, VD, SGD, CT and AC), the pelvic ratio (SGD/TD), PIA and POA. In other words, the low CV and SGD values revealed a significant reduction of the pelvic height; the abnormal SGD/TD ratio indicated a modification of the pelvis proportions; and the abnormally low CT value reflected the rotation of the left hemipelvis towards the inside (Fig 1A).

**Table 4.**
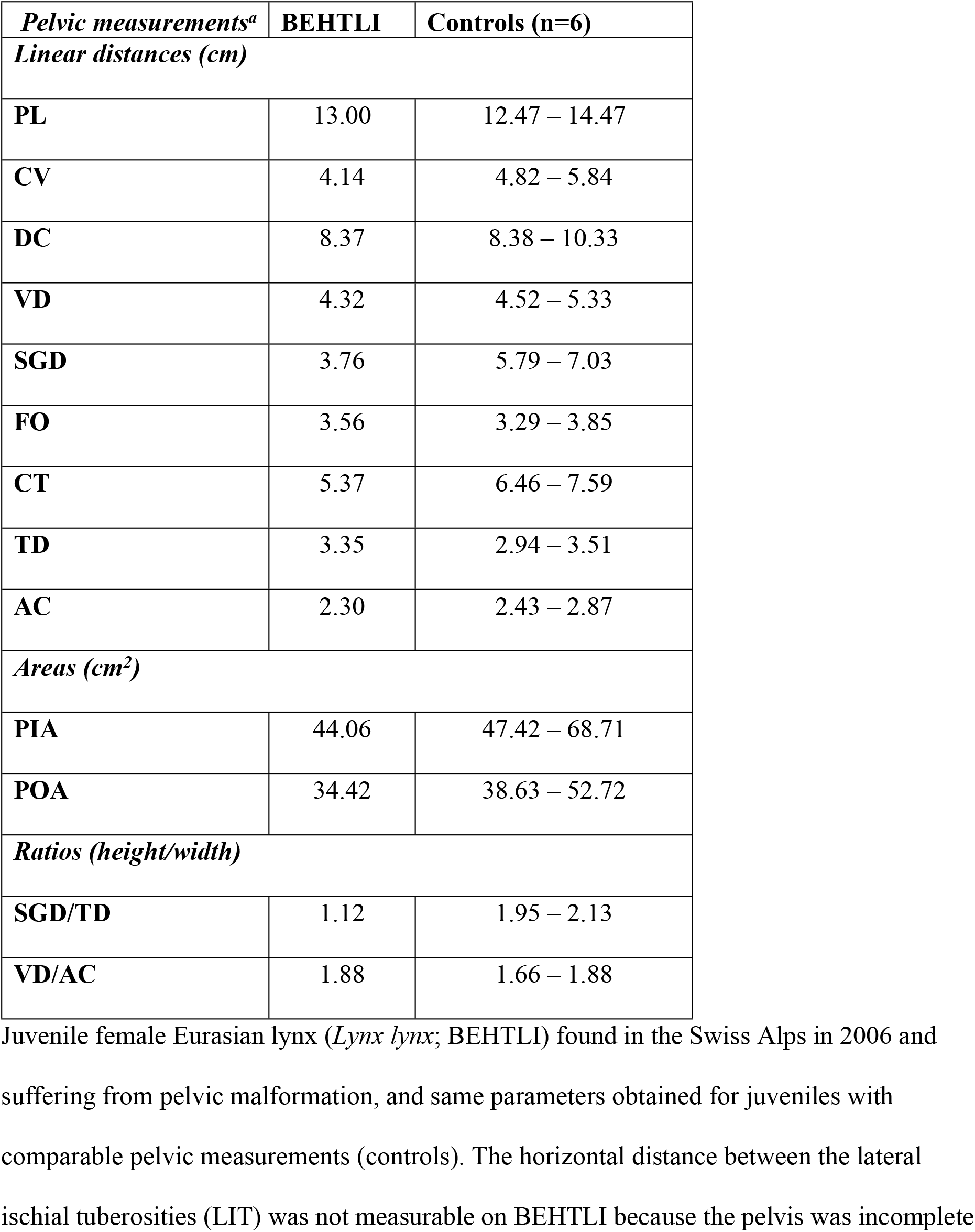
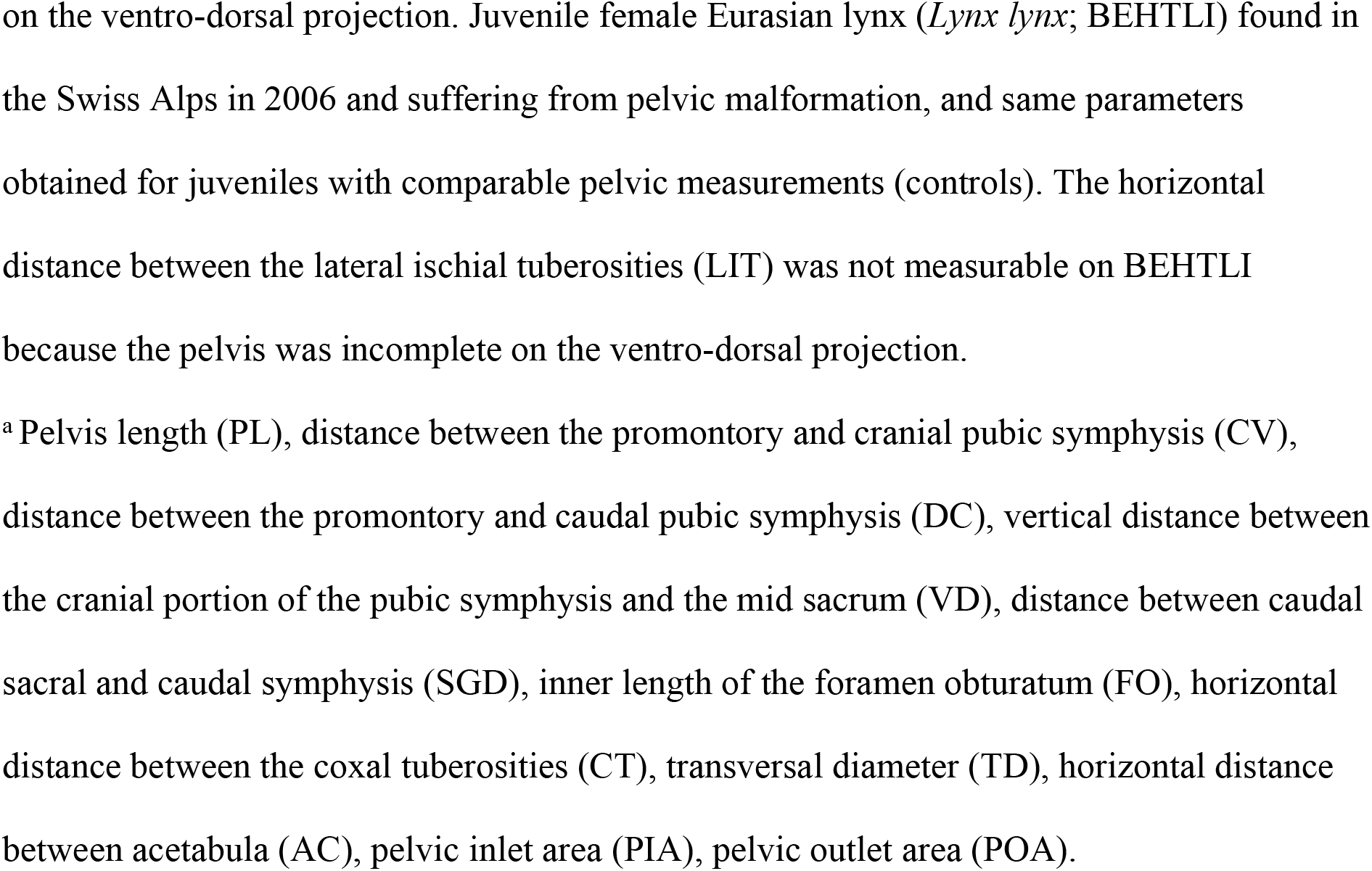
Parameters of an abnormal pelvis.

| *Pelvic measurements<sup>a</sup>* | **BEHTLI** | **Controls (n=6)** |
| --- | --- | --- |
| *Linear distances (cm)* |  |  |
| **PL** | 13.00 | 12.47 – 14.47 |
| **CV** | 4.14 | 4.82 – 5.84 |
| **DC** | 8.37 | 8.38 – 10.33 |
| **VD** | 4.32 | 4.52 – 5.33 |
| **SGD** | 3.76 | 5.79 – 7.03 |
| **FO** | 3.56 | 3.29 – 3.85 |
| **CT** | 5.37 | 6.46 – 7.59 |
| **TD** | 3.35 | 2.94 – 3.51 |
| **AC** | 2.30 | 2.43 – 2.87 |
| *Areas (cm<sup>2</sup>)* |  |  |
| **PIA** | 44.06 | 47.42 – 68.71 |
| **POA** | 34.42 | 38.63 – 52.72 |
| *Ratios (height/width)* |  |  |
| **SGD/TD** | 1.12 | 1.95 – 2.13 |
| **VD/AC** | 1.88 | 1.66 – 1.88 |

Juvenile female Eurasian lynx (*Lynx lynx*; BEHTLI) found in the Swiss Alps in 2006 and suffering from pelvic malformation, and same parameters obtained for juveniles with comparable pelvic measurements (controls). The horizontal distance between the lateral ischial tuberosities (LIT) was not measurable on BEHTLI because the pelvis was incomplete
on the ventro-dorsal projection. Juvenile female Eurasian lynx (*Lynx lynx*; BEHTLI) found in the Swiss Alps in 2006 and suffering from pelvic malformation, and same parameters obtained for juveniles with comparable pelvic measurements (controls). The horizontal distance between the lateral ischial tuberosities (LIT) was not measurable on BEHTLI because the pelvis was incomplete on the ventro-dorsal projection.
<sup>a</sup> Pelvis length (PL), distance between the promontory and cranial pubic symphysis (CV), distance between the promontory and caudal pubic symphysis (DC), vertical distance between the cranial portion of the pubic symphysis and the mid sacrum (VD), distance between caudal sacral and caudal symphysis (SGD), inner length of the foramen obturatum (FO), horizontal distance between the coxal tuberosities (CT), transversal diameter (TD), horizontal distance between acetabula (AC), pelvic inlet area (PIA), pelvic outlet area (POA).

## Discussion

Pelvimetry studies have been performed in a range of animal species but no information was formerly available for Eurasian lynx. A comparison with data on Canada lynx (*L. canadensis*) and Iberian lynx (*L. pardinus*; (18,19) was hampered by a range of factors: species measurements for these studies were taken on prepared bones and not on radiographs, these two lynx species are smaller in size than the Eurasian lynx, part of the parameters differed among studies, and data on *L. pardinus* were taken only on one adult specimen of unknown sex. Future studies on lynx may consider comparing radiological data with direct bone measurements. The latter can be performed post-mortem only but can be done on museum specimens, does not require radiographic equipment and is associated with lower costs, which may permit access to larger sample sizes in more lynx populations.

We made our comparisons with studies in domestic cats (5,8) and dogs (9,10) since the applied methods were the same. Data on pelvimetry remain scarce even in domestic animals and especially in males, because studies mostly focus on the contribution of female pelvic size to rearing performance (9,11,12); however, depending on their pelvic shape, animal species can be classified as dolicopelvic, platypelvic or mesatipelvic. In Eurasian lynx, the values of the pelvic height/width ratios (SGD/TD and VD/AC) were close to one, corresponding to a nearly circular pelvis shape or mesatipelvis for both sexes. Thus, the Eurasian lynx can be classified as a mesatipelvic species, similarly to domestic cats (8).

### Differences between populations

The pelvic measurements and calculated ratios did not significantly differ between the two populations, i.e. pelvis shape was similar for Alpine and Jura lynx. Pelvis size of Alpine lynx seemed to be generally larger but a potential body size difference between the two populations could not be assessed in the absence of additional morphological data, and the statistical analysis was limited by the sample sizes obtained for each sex/age category.

### Growth-related changes

Both pelvic areas and almost all linear pelvic measurements significantly increased with age, as expected from animals in growth. By contrast, the height/width ratios decreased with age in females and only partly in males (Table 2), likely because the pelvis of females increases more markedly in width than in length during the growth period. Similar observations were reported for domestic cats (5).

### Sexual dimorphism

A significant difference between male and female juveniles was detected only for VD but comparisons among lynx <1 year were subject to errors due to rapid body growth and the resulting wide range of pelvic measurements in this age class. This dataset is expected to be particularly influenced by the sample size and possible uneven age distribution for each sex. Similarly, the lack of statistical difference between sexes in the subadult class may have been due to growth and low sample sizes; alternatively, obvious body size differences between sexes may be absent in lynx before the adult age. In domestic cats, only few sex-related differences were observed in 1 year old animals, sexual dimorphism becoming evident only when cats reach 2-3 years (5).

In adult lynx, the pelvis length, CV, DC, VD and SGD were significantly larger in males, in agreement with observations in adult domestic cats (5). Male lynx also had larger PIAs and POAs; this difference was not significant but considering the generally larger body size of males, PIA and POA may be proportionally larger in female lynx. Mesaticephalic female domestic cats have a significantly larger PIAs and POAs than males, and PIA and POA represent important parameters to describe the pelvic space for foetus passage in cat reproduction (5,8). Sex-related body size difference are likely less marked in domestic cats than in Eurasian lynx, potentially explaining the lack of significant difference in lynx. Similarly, two parameters reflecting the pelvic width (TD and AC) were higher in female than male lynx but this was not statistically significant; considering that the pelvis of females is generally smaller than the pelvis of males, proportionally speaking, the larger values in females are likely relevant. In agreement with this, the SGD/TD ratio was significantly larger in males, confirming that at the adult stage lynx females have a different pelvic shape than males, obviously to fulfil the physiological requirements of parturition. Thus, these values need to be considered in case of a rectal obstipation or dystocia in lynx.

Sexual dimorphism of the pelvic bone has been described in humans (20), non-human primates (21), Retriever dogs (22) and domestic cats (8), whereas it is not present in the German shepherd dog (10). This dimorphism can be characterized by a difference in pelvic size and/or pelvic conformation, and if these differences are marked, they can contribute to the sex determination of an individual (20,22). In Eurasian lynx, Canada lynx and domestic cat (5,8,18), sexual dimorphism of the pelvis is characterized by a larger pelvis size in males, in agreement with their larger body size (17,18,23), and by a different pelvic conformation, the pelvis of females being wider.

### Outliers

Outliers which may hint at abnormal morphological features in single individuals were detected twice. Within the juvenile age class, the 10 months old male W09/1096/058 showed CT and TD values clearly above the range of the other lynx, while all other linear measurements, ratios and areas were within the same range. However, this male was the individual with the largest values within the juveniles for all linear measurements except AC (a juvenile female from the Alps had a higher value), and its pelvic measurements, ratios and areas were within the range of subadults, except for the pelvis length, which was slightly smaller (14.47 cm; subadult male range: 14.53–16.77). Since its age was determined based on field data, it excludes a wrong classification. Therefore, despite its age, for an unknown reason this lynx already possessed the morphological conformation of a young subadult specimen and did not have a malformed pelvis.

The other animal with a value outside the range obtained for the same age/sex class was the adult male SIBO, which had a smaller VD. All others pelvic measurements were situated on the verge of the minimal range, except for the pelvis length (16.40 cm), which was close to the mean (17.30 cm ± 0.62, adult male range 14.07–17.97 cm). A comparison with another adult male (JURO, captured in the field in December 2011) with almost the same pelvic length (16.43 cm) suggested that SIBO may have had a pelvic malformation, characterized by abnormally small pelvic measurements for its pelvic length: the PIA and POA of SIBO were 61.93 cm^2^ and 56.08 cm^2^, respectively, while for JURO these values were 69.40 cm^2^ and 72.28 cm^2^. Thus, SIBO presented similar morphological pelvic changes as the juvenile female with the severe malformation (BEHTLI).

BEHTLI’s data indicated a pathological reduction of the pelvic canal. This supports the hypothesis that this malformation may have predisposed to tenesmus and finally rectal perforation due to a foreign body in the feces. Overall, we detected three lynx from the Alps with similar abnormalities, including two orphans in a rehabilitation centre and a radio-collared adult killed by poachers. In our study, these animals represent exceptions but given the relatively high proportion of specimens which could not be measured (38% missing data), the occurrence of more lynx with malformations cannot be excluded. Furthermore, lynx with a malformed pelvis are more likely to die of obstipation, peritonitis or dystocia early in life and may not be found. Carcasses of lynx which are not fitted with a radio-collar and die of non-anthropogenic causes are indeed less likely to be discovered (16).

## Conclusions

The Eurasian lynx can be classified as a mesatipelvic species with sexual dimorphism of the pelvis bone. We identified one adult individual with a value outside the calculated reference range, raising the suspicion of a possible pathological morphology of the pelvic canal. In the juvenile lynx with a pelvis malformation detected at morphological examination, pelvimetry represented an objective tool allowing a quantification of the deviation from normality. Overall, radiographic examinations of the pelvis have proven to be a useful component of the long-term health monitoring of lynx. From a technical point of view, a minimum of two orthogonal projections, including a latero-lateral and a ventro-dorsal projection, are required for pelvimetric analyses. Considering the infrastructure, costs and efforts necessary for radiology and pelvimetry, long-term routine examinations may not be feasible in practice, but these investigations should be considered at least for all animals with a health disorder possibly related to the pelvis conformation, to compare the measurements obtained with the baseline data presented here. Furthermore, the collected data could be useful for estimating age and sex in skeletal remains.

## Acknowledgements

Many thanks go to the hunting authorities, game wardens and biologists of the KORA who organized capture attempts and submitted lynx for post-mortem examination. We also acknowledge the collaborators of the FIWI and of the Department of Clinical Radiology from the Small Animal Clinic for their contributions to lynx pathological and radiographic examinations, respectively.

## Supporting information captions

**S1 Table. P-values of the statistical tests.**

**S1A Table.** P-values of the MANOVA for the effect of age, sex and their interaction on the pelvic measurements of Eurasian lynx (*Lynx lynx*) from Switzerland (1997-2015). Except for the areas PIA and POA (cm^2^) and the two ratios, measurements are given in cm. P-values >0.05 are not shown. An effect of the sex alone, population alone, and interactions population*age, population*sex and population*age*sex was not detected for any of the parameters.

**S2B Table.** P-values of the unpaired Student t test or Mann-Whitney U test applied to pelvic measurements to compare sexes within the same age group of Eurasian lynx (*Lynx lynx*) from Switzerland, 1997-2015. Except for the ratio, measurements are given in cm. Parameters for which no statistical differences and single values ≥0.05 are not shown. No difference was found in subadult lynx.

## References

1. Chapron G, Kaczensky P, Linnell JDC, von Arx M, Huber D, Andrén H, et al. Recovery of large carnivores in Europe’s modern human-dominated landscapes. Science. 2014 Dec 19;346(6216):1517–9.

2. Breitenmoser-Würsten C, Obexer-Ruf G. Population and conservation genetics of two reintroduced lynx *(Lynx lynx)* populations in Switzerland – a molecular evaluation 30 years after translocation. Environ Encount. 2003;58:51–5.

3. Ryser-Degiorgis M-P, Breitenmoser-Würsten C, Meli M, Breitenmoser U. Health surveillance as an important tool in wildlife conservation experiences with the Eurasian lynx. In: Proceedings of the 62th International Conference of the Wildlife DIsease Association. Knoxville, Tennesse, USA; 2013. p. Abstract 46.

4. Schrader SC. Pelvic osteotomy as a treatment for obstipation in cats with acquired stenosis of the pelvic canal: six cases (1978-1989). J Am Vet Med Assoc. 1992 Jan 15;200(2):208–13.

5. Celimli N, Seyrek Intas D, Yilmazbas G, Seyrek Intas K, Keskin A, Kumru IH, et al. Radiographic pelvimetry and evaluation of radiographic findings of the pelvis in cats with dystocia. Tierärztl Prax Kleintiere. 2008;4:277–84.

6. Batista-Arteaga M, Santana M, Lozano O, Méndez J, Quesada O, Arbelo M, et al. Medical and surgical management of a dystocia because of foetopelvic disproportion in an African lioness *(Panthera leo)*. Reprod Domest Anim Zuchthyg. 2011 Apr;46(2):362–5.

7. Linde-Forsberg C. Pelvimetry to diagnose dystocia in the bitch. In: Proceedings of the 27th World Small Animal Veterinary Association World Congress. Granada, Spain; 2002. p. 591.

8. Monteiro CLB, Campos AIM, Madeira VLH, Silva HVR, Freire LMP, Pinto JN, et al. Pelvic differences between brachycephalic and mesaticephalic cats and indirect pelvimetry assessment. Vet Rec. 2013 Jan 5;172(1):16.

9. Eneroth A, Linde-Forsberg C, Uhlhorn M, Hall M. Radiographic pelvimetry for assessment of dystocia in bitches: a clinical study in two terrier breeds. J Small Anim Pract. 1999 Jun;40(6):257–64.

10. Ocal M, Dabanoglu I, Kara ME, Turan E. Computed tomographic pelvimetry in German shepherd dogs. 2003;110(1):17–20.

11. Van Donkersgoed J. A critical analysis of pelvic measurements and dystocia in beef heifers. Compend Contin Educ Pract Vet. 1992;14:405–7.

12. Cloete SW, Haughey KG. Radiographic pelvimetry for the estimation of pelvic dimensions in Merino, Dormer and S A mutton Merino ewes. J S Afr Vet Assoc. 1990 Jun;61(2):55–8.

13. Malik MR, Crao K, Taluja JS, Shrivastava AM. Length and girth as an index to surface pelvimetry in buffalo. Indian J Anim Sci. 1990;60:1200–2.

14. Vogt K, Zimmermann F, Kölliker M, Breitenmoser U. Scent-marking behaviour and social dynamics in a wild population of Eurasian lynx *Lynx lynx*. Behav Processes. 2014 Jul;106:98–106.

15. Vogt K, Hofer E, Ryser A, Kölliker M, Breitenmoser U. Is there a trade-off between scent marking and hunting behaviour in a stalking predator, the Eurasian lynx, *Lynx lynx*? Anim Behav. 2016 Jul 1;117:59–68.

16. Schmidt-Posthaus H, Breitenmoser-Würsten C, Posthaus H, Bacciarini L, Breitenmoser U. Causes of mortality in reintroduced Eurasian lynx in Switzerland. J Wildl Dis. 2002 Jan;38(1):84–92.

17. Yom-Tov Y, Kjellander P, Yom-Tov S, Mortensen P, Andrén H. Body size in the Eurasian lynx in Sweden: dependence on prey availability. Polar Biol. 2010 Apr 1;33(4):505–13.

18. Garciá-Perea R. Variabilidad morfologica del genero *Lynx* Kerr, 1792 (Carnivora: Felidae). [Madrid, Spain]: Universidad Complutense de Madrid; 1990.

19. Gálvez Prada F, Serrano León JP, Sañudo Franquelo B, Beltrán Gala JF. Digital atlas of the skeleton of *Lynx pardinus* [Internet]. Sevilla, Spain: BioScripts & Universidad de Sevilla; 2013 [cited 2015 Nov 17]. Available from: http://thevirtualmuseumoflife.com/lynx/index-en.php?bone=097

20. Gonzalez PN, Bernal V, Perez SI. Geometric morphometric approach to sex estimation of human pelvis. Forensic Sci Int. 2009 Aug 10;189(1–3):68–74.

21. Leutenegger W, Larson S. Sexual dimorphism in the postcranial skeleton of New World primates. Folia Primatol Int J Primatol. 1985;44(2):82–95.

22. Nganvongpanit K, Pitakarnnop T, Buddhachat K, Phatsara M. Gender-Related Differences in Pelvic Morphometrics of the Retriever Dog Breed. Anat Histol Embryol. 2017 Feb;46(1):51–7.

23. Pontier D, Fromont E, Courchamp F, Artois M, Yoccoz NG. Retroviruses and sexual size dimorphism in domestic cats (Felis catus L.). Proc Biol Sci. 1998 Feb 7;265(1392):167–73.

